# HiReS: A Method for Automated Morphometric Trait Extraction from High-Resolution Plankton Images

**DOI:** 10.64898/2026.04.11.717915

**Authors:** Stylianos Mavrianos, Sven Teurlincx, Steven AJ Declerck, Kathrin A Otte

**Affiliations:** Institute of Animal Cell and Systems Biology, Department of Biology, University of Hamburg,Hamburg, Germany; Department of Aquatic Ecology, Netherlands Institute of Ecology (NIOO-KNAW), Wageningen, The Netherlands; Laboratory of Aquatic Ecology, Evolution and Conservation, Department of Biology, KULeuven, Leuven, Belgium

**Keywords:** *Daphnia*, High-resolution image analysis, Instance segmentation, Machine learning, Morphometric trait extraction, Plankton ecology

## Abstract

Trait-based analyses in plankton ecology require measurements from large numbers of individuals, yet morphometric data are typically collected manually from small subsets. Although deep learning methods enable automated detection and segmentation, extracting quantitative trait data from full-resolution images remains challenging due to memory limitations. We present HiReS (High-Resolution Segmentation), an open-source workflow for automated morphometric trait extraction from large plankton images. HiReS partitions images into overlapping chunks, performs YOLO-based instance segmentation, reconstructs polygon annotations in full-image space, removes truncated and duplicate detections, and computes geometric descriptors. We evaluated the workflow using manually annotated and automated segmentations of *Daphnia pulex, Daphnia galeata*, and *Simocephalus vetulus*. Automated measurements reproduced the structure of manual trait distributions and showed strong agreement at both sample and individual levels. A consistent positive bias was observed, reflecting a multiplicative scaling offset rather than distortion of relative trait structure. Subsampling analyses further showed that model-derived medians can outperform manual estimates at low sampling depths. HiReS provides a reproducible and computationally efficient framework for extracting morphometric traits from full-resolution plankton images.

## 1. Introduction

Ecology aims to understand how organismal characteristics mediate interactions with the environment and scale to shape community structure and ecosystem functioning. The ultimate goal is to predict how ecosystems respond to environmental pressures. Functional trait-based approaches (FTBAs) have emerged as a unifying framework to address this objective by linking measurable phenotypic properties to ecological performance, demographic processes, and biogeochemical dynamics (Kremer et al., 2017; Martini et al., 2021). Rather than focusing solely on taxonomic identity, trait-based ecology emphasizes quantitative, mechanistic descriptors that enable prediction of ecosystem responses to environmental drivers. In planktonic systems, morphometric traits are particularly informative. Body size governs metabolic scaling relationships and energy transfer efficiency, while shape and structural complexity influence feeding performance, predator avoidance, and sinking rates (Banse, 1976; Woodward et al., 2010; Yvon-Durocher et al., 2012). The full realization of trait-based aquatic ecology depends on the development of standardized, high-throughput trait extraction tools (Kremer et al., 2017). Likewise, achieving cross-system comparability and predictive trait-based modelling requires reproducible and scalable measurement pipelines (Martini et al., 2021).

For decades, assessments of mesocosm and laboratory samples have relied on manual microscopy, where organisms are visually identified, counted, and measured (Adoteye et al., 2015; McKnight et al., 2023). While this approach has underpinned much of our current ecological knowledge, it is inherently time-consuming, labor-intensive, and limits the scale of ecological inquiry. Moreover, manual identification and measurement are subject to substantial operator-dependent variation. Differences in experience and judgment can introduce inconsistencies that compromise the reproducibility and comparability of long-term and collaborative datasets (Culverhouse et al., 2014; Lindenmayer & Likens, 2010).

Advances in imaging technologies, including ZooScan (Gorsky et al., 2010) and FlowCAM (Sieracki et al., 1998) systems, have enabled the rapid acquisition of high-resolution images containing thousands of planktonic individuals per sample. These platforms have substantially increased throughput and improved standardization in plankton monitoring. However, the extraction of biologically meaningful traits from the resulting large image datasets has not kept pace with image acquisition. In practice, researchers still frequently measure only a small subset of individuals and extrapolate results to the broader community (Lindenmayer & Likens, 2010; Milbrink & Bengtsson, 1991; Orenstein et al., 2022). As a result, much of the information contained in full-resolution scans remains underutilized.

This imbalance between image acquisition and data extraction has led to growing interest in automated analysis methods. Advances in deep learning, particularly in computer vision methods, have demonstrated strong potential in ecological image analysis (Bachimanchi et al., 2024; Chen et al., 2025; Karatzas et al., 2020; Kim et al., 2023; Kyathanahally et al., 2021; Lumini et al., 2023; Lumini & Nanni, 2019; Ma et al., 2024; Rutter et al., 2024). These approaches enable automated identification and counting of planktonic organisms, but typically do not extract quantitative morphometric traits. Among available architectures, YOLO (You Only Look Once)–based instance segmentation models are particularly suited to high-throughput ecological applications (Jocher & Qiu, 2024). YOLO performs object localization and segmentation in a single forward pass, allowing efficient inference even in images containing large numbers of individuals (Sapkota et al., 2024). YOLO instance segmentation models generate polygon-based masks that delineate the full spatial extent of each organism. This capability is essential for trait-based analyses, as geometric descriptors derived from complete object outlines capture biologically meaningful variation in body size and shape. Recently, YOLO-based instance segmentation frameworks have begun to emerge for planktonic organisms, combining base segmentation models with transfer learning approaches for species-level identification (Mavrianos et al., under review) .

Despite these developments, a critical computational barrier remains. High-resolution scans frequently exceed 10,000 x 10,000 pixels, generating images that are too large to be processed in a single forward pass by standard computer. During inference, the entire image must reside in CPU memory (Sapkota et al., 2024), and because memory usage scales with image resolution and feature-map depth, high-resolution ecological images often exceed available hardware capacity, resulting in memory errors or prohibitive computational demands. To address this limitation, slicing-based inference strategies have been developed that divide large images into smaller chunks and subsequently merge their predictions. For example, Slicing-Aided Hyper Inference (SAHI) applies this strategy to improve detection of small objects in large images (Adiwijaya et al., 2024). Similarly, segmentation-based pipelines have been developed for ecological monitoring, such as the flatbug framework, which uses instance segmentation to detect terrestrial arthropods across diverse imaging systems (Svenning et al., 2026). While these approaches improve automated organism detection in large images, their outputs primarily consist of detected or segmented individuals rather than quantitative morphometric measurements.

In this study, we present an open-source workflow designed to bridge the gap between high-resolution image acquisition and quantitative trait analysis in plankton research. Our framework integrates HiReS (High-Resolution Segmentation), a chunk-based segmentation strategy that overcomes the memory limitations associated with processing full-resolution images, with the direct extraction of morphometricc traits from polygon-based instance outlines. Building on our previous work in Mavrianos et al., under review (DaphnAI framework), which focused on accurate organism detection and taxonomic classification, the present study addresses the subsequent challenge of processing segmentation outputs at the scale of full-resolution ecological images and converting reconstructed outlines into quantitative trait data. To validate its performance, we compared automated trait measurements against manually derived ground truth data, focusing on key shape descriptors such as surface area, width, and length. From these segmentations, we compute quantitative metrics that enable high-throughput, multivariate trait analyses at ecologically relevant scales. Importantly, the proposed approach is not tied to a specific image acquisition platform or segmentation model, allowing it to be flexibly applied across different imaging systems and YOLO-based model architectures.

## 2. Materials and Procedures

### 2.1 Materials

#### 2.1.1 Imaging Data

High-resolution images can be acquired using a flatbed scanner (e.g., Epson Perfection V850 Pro). The sample is placed in an external transparent scanning tray made of flat plastic (127 x 84 x 8 mm), which is positioned directly on the scanner glass. The tray contains a shallow layer of the preservation medium, allowing organisms to settle and distribute within a single imaging plane. This configuration reduces overlap among individuals and helps maintain consistent focus across the image. Prior to scanning, the sample is gently spread as evenly as possible within the tray to minimize aggregation and clustering. This approach allows entire samples to be digitized in a single scan and provides a cost-effective alternative for image acquisition. Although this study uses flatbed scanning, the image acquisition step is not restricted to this approach. The workflow is also compatible with images generated by other systems commonly used in plankton research, including platforms such as FlowCAM or ZooScan (Gorsky et al., 2010; Siekiera et al., 2025), as well as other imaging setups that produce sufficiently high-resolution images with clear object boundaries.

#### 2.1.2 Segmentation Models

To perform instance segmentation on the high-resolution images, the workflow requires a deep-learning model capable of detecting and segmenting objects of interest. The workflow is fully class-agnostic, meaning that it functions independently of the specific classes the model was trained on and accepts any YOLO segmentation model that outputs polygon-based annotations in the standard YOLO format. When the study (e.g., species composition and imaging conditions) closely matches an existing dataset (i.e., experiments with similar species compositions), it is advisable to use pretrained community models. This applies to both controlled experimental datasets and field samples. These models can be downloaded and used directly by supplying the corresponding .*pt* weight file to the segmentation step, allowing immediate analysis without the need for custom annotation.

In cases where no appropriate model exists (such as experiments involving novel species, unique imaging conditions, or specific structural targets) users may train their own model. For this purpose, the workflow provides a chunking command that converts full-resolution images into chunks suitable for manual annotation. These chunks can be imported into annotation tools such as SegmentME (https://segmentme.streamlit.app/), a desktop-based interactive image annotation application built with PyQt6 that enables semi-automatic segmentation and manual refinement of object outlines. Importantly, chunking is not limited to model training or annotation but is also a core component of the inference pipeline, including when pretrained models are used, as it enables processing of full-resolution images that would otherwise exceed memory constraints. General principles and workflows for training deep learning-based computer vision models in ecological contexts are discussed in (Blair et al., 2024; Chen et al., 2025; Christin et al., 2019; Cole et al., 2023).

#### 2.1.3 Software and Hardware

HiReS is an open-source computational python library enabling efficient processing of large images in a reproducible manner. Its primary function is to overcome memory limitations associated with extracting FTBAs from full-resolution images by implementing a standardized chunk-based segmentation strategy. HiReS is implemented as both a command-line interface and a Python API and can be downloaded from PyPI (https://pypi.org/project/HiReSeg/) through pip or can be cloned from our GitHub repository (https://github.com/StevetheGreek97/HiReS), enabling high-resolution image segmentation and quantitative trait extraction within reproducible analytical workflow.

The library provides three primary operations. The *chunk* command partitions large images or directories with images into overlapping chunks with user-defined dimensions and overlap, facilitating both efficient processing and manual annotation by breaking down high-resolution images into manageable chunks. The *run* command executes the complete high-resolution segmentation pipeline. In this mode, user-supplied images are combined with a specified YOLO-based instance segmentation model, either pretrained or custom-trained. The pipeline then performs inference on chunks, integrates model predictions into a unified full-image representation, and converts reconstructed object outlines into quantitative measurements of individual organisms. This pipeline therefore links raw image data to structured morphometric datasets within a single reproducible process. Finally, the *plot* command generates visualization overlays by combining source images with YOLO-format annotations, supporting result verification and preparation of publication-quality figures.

In addition to flexibility in image acquisition, the computational requirements of the pipeline are modest. The full workflow, including image chunking, segmentation inference, reconstruction, duplicate removal, and morphometric descriptor extraction, can be executed on standard consumer hardware. In the present study, all processing steps were performed on a laptop equipped with an Intel Core i5 processor (12 CPU cores) and 15 GB RAM, without the use of GPU acceleration. This demonstrates that the pipeline is accessible and can be routinely applied without specialized computing infrastructure.

### 2.2 Procedures

#### 2.2.1 HiReS Pipeline

Once high-resolution images and a YOLO-based instance segmentation model are collected, the hires run command executes the end-to-end processing pipeline. The workflow is designed to enable inference on images that exceed CPU memory limits while preserving geometrically consistent object representations for downstream morphometric analysis (Figure 1).

**Figure 1:**
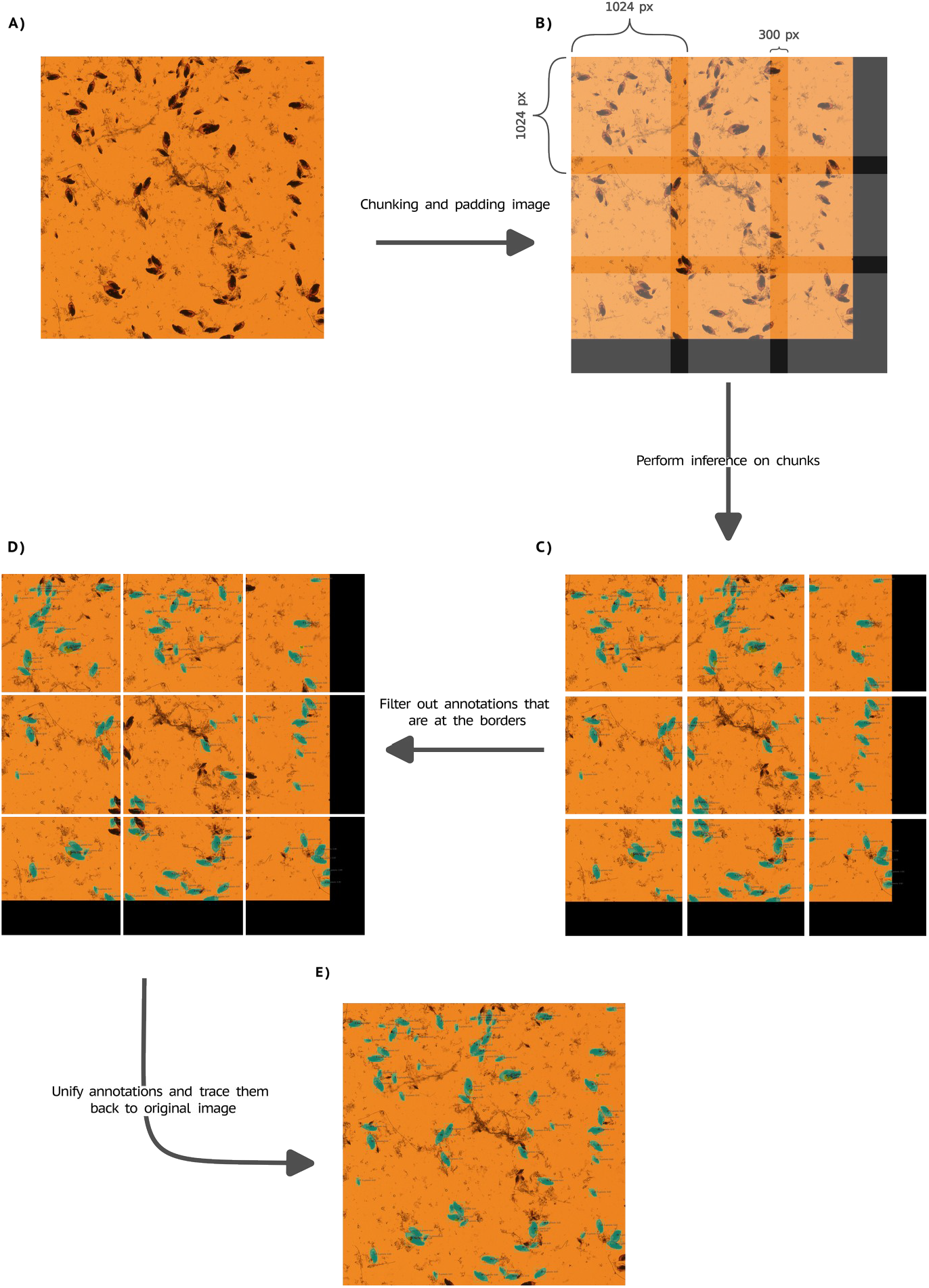
Overview of the pipeline A) Original high-resolution input image (shown here with a 3 3 layout) containing numerous Daphnia individuals and debris. B) Preprocessing step, where the full image is cropped into fixed-size chunks with the specified overlap. C) The segmentation model performs inference independently on each chunk. D) Reconstructed full-image prediction after merging all chunk outputs. Segmentation masks are transformed back to global coordinates and stitched together, while overlapping predictions are resolved using class-agnostic non-maximum suppression to eliminate duplicates and maintain mask consistency.

##### Chunking

The first stage partitions each full-resolution image into a regular grid of overlapping chunks using a sliding-window approach. By default, chunks are 1024 x 1024 pixels with a user-configurable overlap (default: 150 pixels). The stride of the sliding window equals chunk_size plus the overlap, ensuring that adjacent chunks share overlapping regions. Overlap is necessary to guarantee that organisms located near chunk boundaries remain fully visible in at least one chunk and are not represented solely as partial detections. If a chunk extends beyond the image boundary, the area outside the original image is filled with zero-intensity (black) pixels to preserve the fixed chunk size, so that all chunks maintain identical dimensions. This ensures consistent input size during model inference. Each chunk is saved with its top-left pixel offset encoded in the filename ({base}_{x}_{y}.png), where x and y denote absolute pixel coordinates in the original image. These offsets are later used to reconstruct predictions into full-image space (Figure 1a & b.

##### Inference & Filtering

Then, each chunk is processed independently using the specified YOLO instance segmentation model. Predictions are written in YOLO segmentation format, consisting of class identifiers followed by polygon vertices expressed in chunk-normalized coordinates (values between 0 and 1 relative to chunk dimensions). Because objects intersecting chunk boundaries may produce truncated polygons, HiReS applies a geometry-based edge filter at the chunk level. Polygons are retained only if all vertices lie within a slightly inset unit square defined by a small inward buffer (edge threshold; default 1 x 10^-5^). This removes detections that touch or cross chunk borders and therefore lack biological interpretability. For retained objects, polygon vertices are transformed from chunk-local to global image coordinates. This is performed in three steps: i) Denormalization, chunk-normalized coordinates are converted to pixel units using chunk dimensions. ii) Offset correction, the chunk’s horizontal and vertical offsets (parsed from the filename) are added to each vertex, shifting the polygon into full-image pixel space. iii) Global normalization,coordinates are re-normalized relative to full-image dimensions. Finally, transformed polygons from all chunks are merged into a unified collection representing the entire image. A second edge filter is applied at the full-image level to remove any residual border-touching detections from the padded images (Figure 1c & d).

##### Merging

Because overlapping chunks may yield multiple detections of the same instance (e.g., organism), HiReS applies polygon-level non-maximum suppression (NMS) to the unified annotation set. A spatial index (STRtree; (Leutenegger et al., 1997) is constructed to efficiently identify polygon pairs with overlapping bounding boxes. Intersection-over-union (IoU) is then computed for candidate pairs on polygon level. Polygons exceeding a user-defined IoU threshold (default: 0.7) are considered duplicates, and only the highest-confidence detection is retained. This procedure ensures that each organism is represented by a single, geometrically consistent polygon while maintaining computational scalability for large images (Figure 1e).

##### Output

The final component of the pipeline provides a structured file for quantitative analysis of segmented objects. After segmentation, merging, and duplicate removal, the unified polygon file is used to calculate a suite of morphometric variables for each object. All morphometric measurements are initially reported in pixel units, reflecting the native resolution of the input images. To obtain biologically meaningful units (e.g. µm), users can convert pixel-based measurements using a calibration factor derived from the imaging setup (e.g. scanner resolution or scale reference), allowing direct translation of measurements into real-world units. The pipeline extracts fundamental descriptors that characterize the physical scale of the organism. The morphometric descriptors extracted here correspond to commonly used size- and shape-based traits in image analysis, including area, body dimensions, and shape complexity metrics, which have been shown to capture ecologically relevant shape variation among organisms (Orenstein et al., 2022). Area (A) is calculated using the Shoelace Formula, which determines the area of a non-self-intersecting polygon by summing the cross-products of its vertex coordinates.

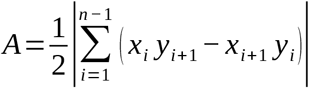

*Equation 1: Quantifies the two-dimensional size of an individual by measuring the surface enclosed by its polygon-based outline. Area provides a direct estimate of body size derived from the full object geometry. Here, (x*_*i*_, *y*_*i*_*)are the Cartesian coordinates of the i-th vertex of the polygon, and n is the total number of vertices. Area provides a direct estimate of body size derived from the full object geometry*.

Perimeter (P) is defined as the cumulative Euclidean distance between consecutive vertices along the polygon boundary.

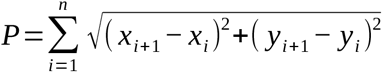

*Equation 2: Measures the total length of the object boundary by summing distances between consecutive points along the polygon outline*. (*x*_*i*_, *y*_*i*_) *and* (*x*_*i*+1_, *y*_*i*+1_) *represent consecutive polygon vertices, and n is the total number of vertices (with the last point connected to the first). Perimeter captures both object size and boundary complexity*.

To determine oriented body dimensions, a Minimum Area Bounding Box is fitted to each polygon. This method identifies the specific orientation that minimizes the area of the enclosing rectangle. The major axis of this box corresponds to the maximum body length, while the minor axis provides the maximum width (SFigure 1; OBB; Oriented Bounding Box).

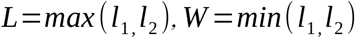

*Equation 3: Characterizes the main body dimensions of an object based on its oriented minimum-area bounding box. Here, l*_1_ *and l*_2_ *are the lengths of the two orthogonal sides of the bounding box. Length (L) represents the longest dimension, while width (W) represents the shortest, providing orientation-independent size estimates*.

Circularity (C) describes the compactness of the object relative to a circle. A value of 1.0 represents a perfect circle, with decreasing values indicating elongated or more complex geometries.

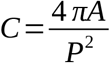

*Equation 4: Measures how closely the shape of an object approximates a circle. A is the area of the object and P is its perimeter. Circularity decreases as objects become more elongated or irregular*.

Convexity (K) describes the smoothness of the object’s boundary. It is calculated as the ratio between the perimeter of the convex hull (Pconvex) and the actual perimeter of the polygon. This ratio identifies high-frequency surface irregularities; a jagged or irregular boundary will yield a lower convexity value than a smooth, continuous edge.

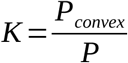

Equation 5: Quantifies boundary smoothness by comparing the perimeter of the object to that of its convex hull. P is the perimeter of the object, and *P*_*convex*_is the perimeter of the convex hull. Values closer to one indicate smoother, more convex outlines, whereas lower values reflect increased boundary irregularity.

Finally, solidity (S) measures the overall density of the shape by comparing the area of the polygon to the area of its convex hull, which is the smallest convex set containing the object. Low solidity values indicate the presence of deep concavities or structural indentations.

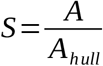

*Equation 6: Describes the degree to which an object fills its convex hull. A is the area of the object, and A*_*hull*_*is the area of its convex hull (the smallest convex polygon enclosing the object). Lower solidity values indicate more concave or irregular shapes*.

The resulting datasets can be exported in CSV format for subsequent statistical analysis or visualization in R, Python, or any other analytical environment.

## 3. Assessment

To evaluate the performance of the workflow, high-resolution images and corresponding manual annotations were taken from the test set of the laboratory Daphniidae Dataset (LDD) described in Mavrianos et al. (under review). This subset comprises nine full-resolution images (three per species: *Daphnia pulex, Daphnia galeata*, and *Simocephalus vetulus*), representing controlled single-species samples typical of laboratory-based ecological experiments. Prediction outputs were generated using the hires run command, which executes the full pipeline, including image chunking, instance segmentation, reconstruction of predictions into full-image space, duplicate removal, and morphometric descriptor extraction. Segmentation was performed using a pretrained YOLO-based model from the DaphnAI framework (Mavrianos et al., under review), a two-stage deep learning approach developed for detection, instance segmentation, and classification of freshwater zooplankton in high-resolution scanner images. In this framework, a general base model is first trained to detect and segment *Daphniidae* individuals independent of species identity and is subsequently fine-tuned for species-level classification. The framework was originally trained and evaluated on both mesocosm and laboratory datasets, representing typical experimental systems used to study zooplankton population dynamics and species composition. While DaphnAI focuses on taxonomic classification and segmentation, the HiReS workflow addresses the subsequent challenge of processing segmentation outputs at the scale of full-resolution images and converting reconstructed outlines into quantitative morphometric descriptors, thereby representing complementary components of an automated plankton image analysis pipeline.

### 3.1 Validation of Automated Morphometric Measurements

To evaluate the outputs of the pipeline, we compared morphometric descriptors derived from automated segmentations with those obtained from manually annotated outlines at three complementary analytical levels: full trait distributions, one-to-one matched object measurements, and sample-level summary statistics. Across all analyses, only true positives were considered to ensure that differences reflect measurement error on correctly detected individuals rather than being confounded by detection errors such as missed or misclassified predictions.

At the distribution level, the pipeline reproduced the overall shape of the manual trait distributions across all three species and all three descriptors (Figure 2). The predicted distributions retained the same broad structure and relative ordering among samples, indicating that the workflow preserved the main biological variation present in the data. However, the predicted distributions were consistently shifted toward larger values. Bland–Altman analysis was used to quantify the agreement between automated and manual measurements at the level of individual matched objects (Bland & Altman, 2007). This method assesses agreement by calculating, for each matched pair, the difference between the two measurement approaches (i.e., ground truth vs prediction) and relating this difference to their mean value. Plotting the difference against the mean allows visualization of both systematic bias and the spread of measurement error across the full range of object sizes. The mean difference represents the average bias between methods, while the dispersion of differences reflects variability and potential heteroscedasticity (i.e. whether error changes with size). Differences were calculated on a log_10_ scale, meaning that they represent multiplicative rather than absolute differences. In addition, fitting a regression line allows assessment of size-dependent bias: a non-zero slope indicates that the magnitude of the difference between methods changes with object size. Specifically, a negative slope indicates that smaller individuals tend to be overestimated more strongly than larger ones, whereas a positive slope would indicate the opposite pattern.

**Figure 2:**
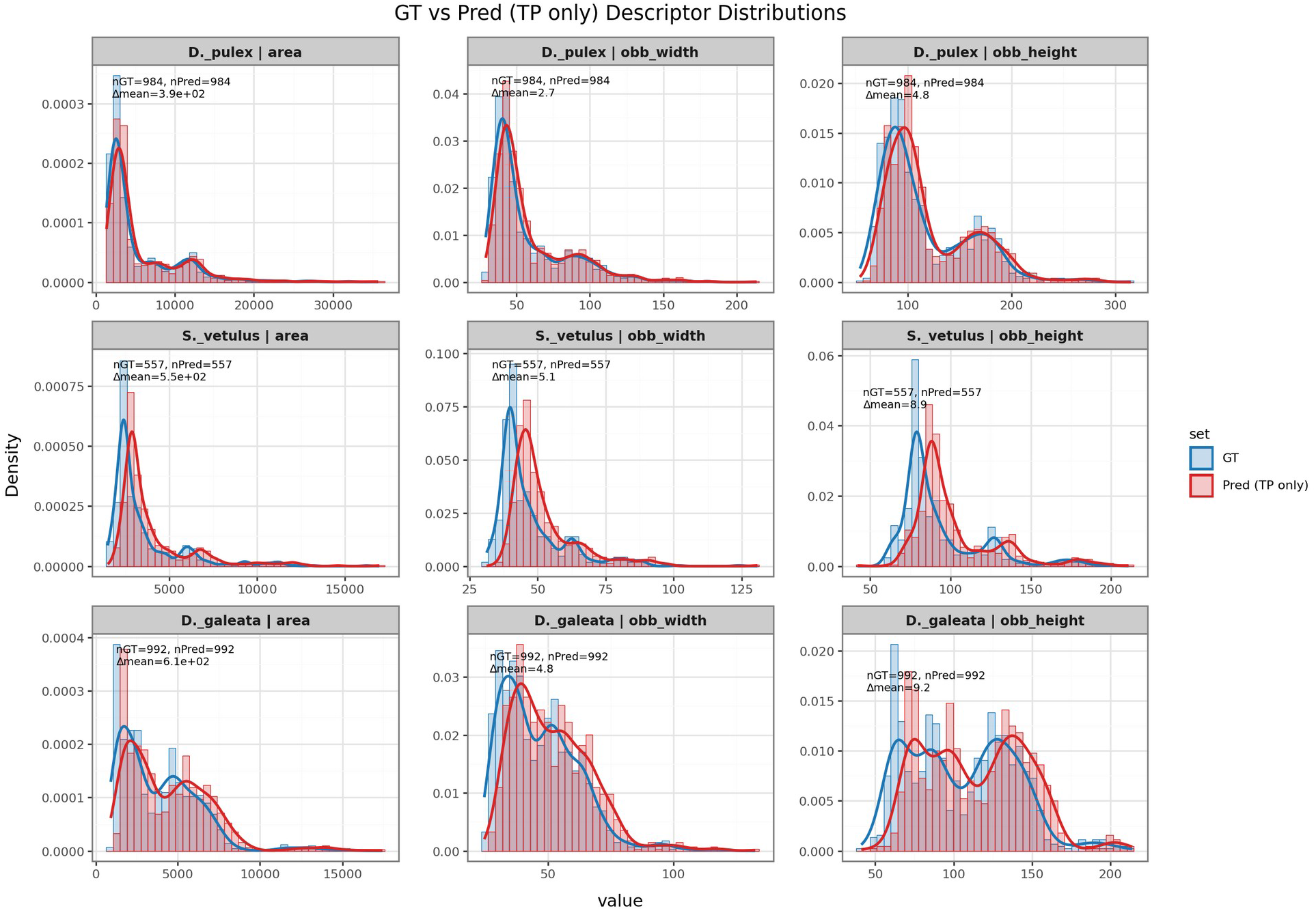
Comparison of ground-truth and automated morphometric descriptor distributions. Density histograms and kernel density estimates showing the distributions of three morphometric descriptors (area, oriented bounding box width, and oriented bounding box height) for D. pulex, D. galeata, and S. vetulus. Blue curves represent descriptors derived from manually annotated outlines (GT), while red curves represent descriptors derived from automated segmentations (Pred). Across species and descriptors, the automated workflow reproduces the overall shape and structure of the manual trait distributions, although predicted values are consistently shifted toward larger measurements. Reported Δmean values represent the mean difference between predicted and manual descriptors (Pred - GT), expressed in pixels, and nGT and nPred denote the number of matched instances included in each comparison. These results demonstrate that automated measurements preserve the relative structure of trait distributions while introducing a systematic positive offset.

The analysis revealed a consistent positive bias of automated measurements relative to the manual reference across all species and descriptors (Figure 3). Mean log-bias values ranged from 0.022 to 0.076 log_10_ units, corresponding to multiplicative offsets of approximately 1.05- to 1.19-fold (5% to 19%). The strongest bias was observed for area in *D. galeata* (0.076; ∼1.19) and *S. vetulus* (0.071; ∼1.18), whereas the smallest deviation occurred for OBB height in D. pulex (0.022; ∼1.05). Overall, OBB height showed the highest agreement with manual measurements across species, indicating that these metric are more robust to segmentation-related biases than area-based descriptors. In addition to this overall bias, we observed a clear size-dependent pattern. Across all species and descriptors, regression slopes were consistently negative (ranging from 0.031 to 0.140), indicating that smaller individuals were more strongly overestimated than larger ones. This trend is visible as a decreasing bias with increasing object size and suggests that measurement error is not constant across the size range.

**Figure 3:**
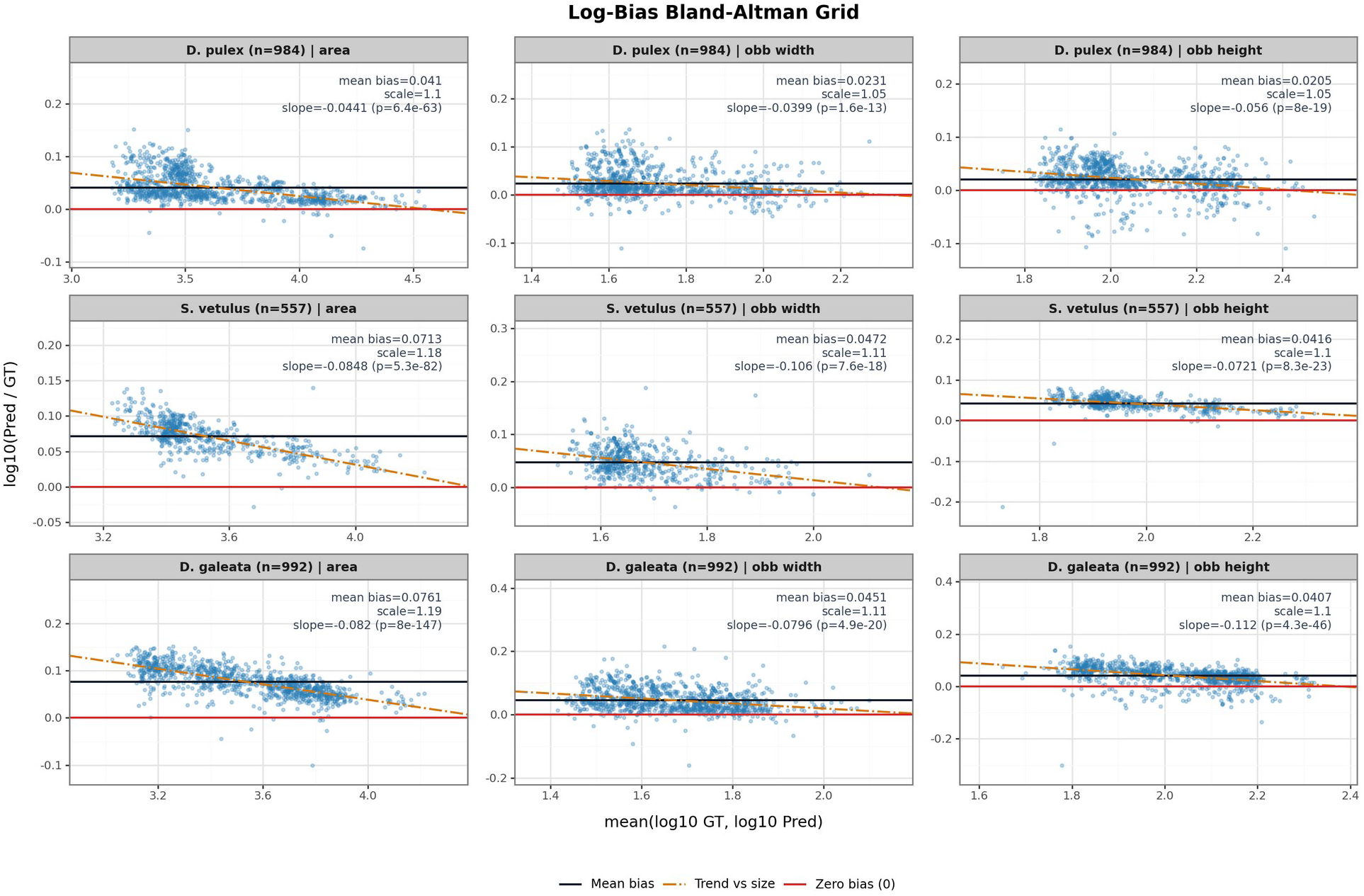
Bland–Altman analysis of automated versus manual morphometric measurements across species and descriptors. Each panel shows the log10(Pred/GT) as indicator of prediction bias plotted against the mean of log-transformed predicted and ground-truth values for area, oriented bounding-box width (OBB width), and oriented bounding-box height (OBB height) in D. pulex, D. galeata, and S. vetulus. Points represent matched measurements for the same biological individuals. The solid black line indicates the trait mean, the dashed orange line shows the linear regression?, and the red horizontal line marks zero bias. Across descriptors and species, automated measurements exhibit a consistent positive bias relative to manual annotations. The negative slopes indicate that this bias decreases with increasing organism size, suggesting stronger overestimation for smaller individuals.

At the sample-summary level (i.e. each image represents a single sample, treatment, or replicate), predicted sample medians remained strongly correlated with correlation coefficients of 0.96 for area, 0.96 for OBB width, and 0.94 for OBB height (Figure 4; row: Original). Despite the positive object-level bias, automated measurements preserved relative differences among samples well, with most points closely aligned along the 1:1 relationship. Absolute deviations between predicted and manual medians were moderate, with mean absolute errors (MAE) of 521 pixels for area, 4.53 pixels for OBB width, and 8.64 pixels for OBB height.

**Figure 4:**
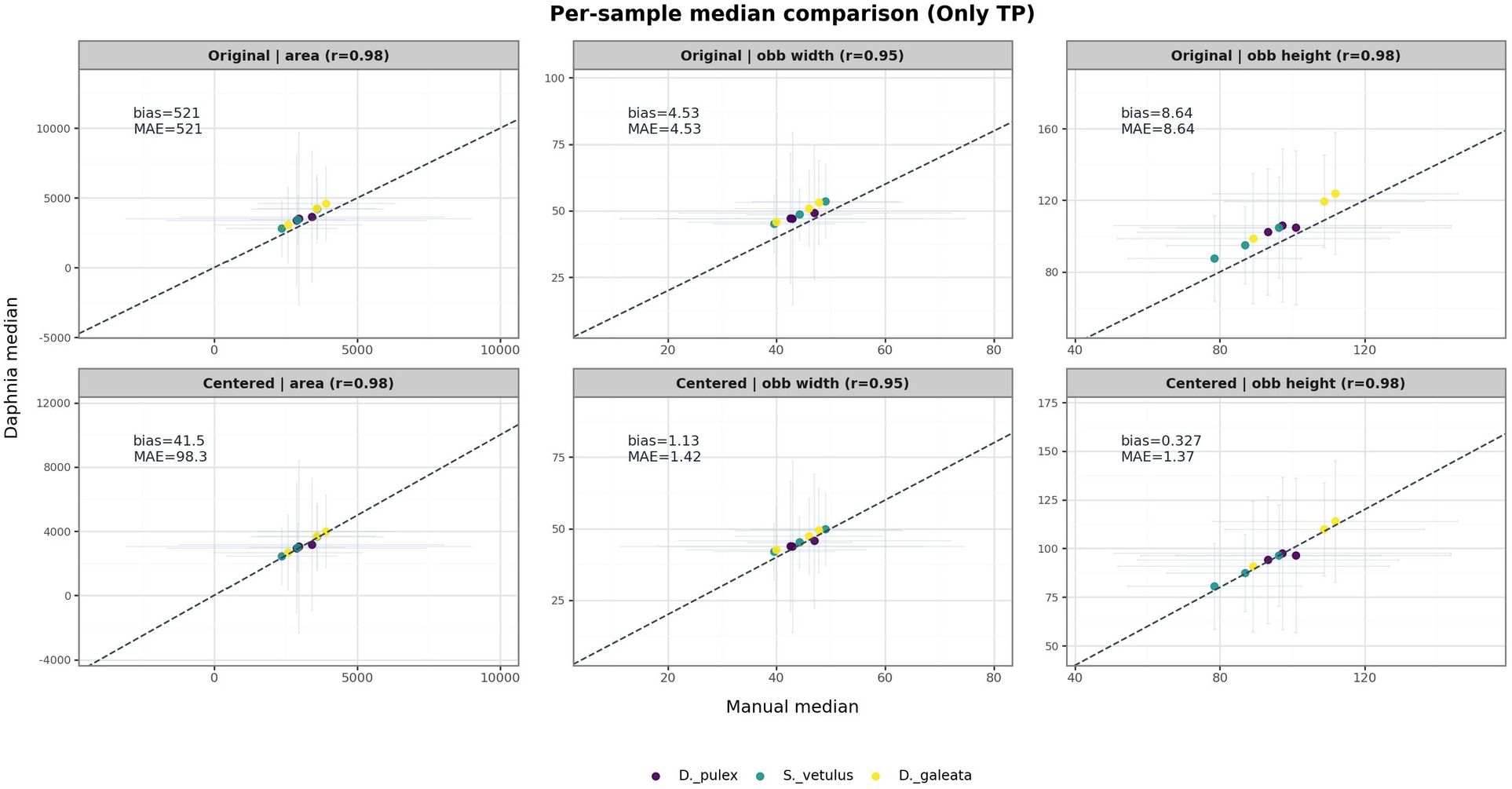
Comparison of sample-level median trait values derived from automated and manual measurements. Each panel shows manually measured medians (x-axis) plotted against prediction-derived medians (y-axis) for three descriptors: area, oriented bounding-box width (OBB width), and oriented bounding-box height (OBB height). Points represent individual samples across the three species (D. pulex, D. galeata, S. vetulus), and the dashed line indicates the identity relationship (1:1). Horizontal and vertical error bars show the spread of the underlying distributions within each sample.In the top row (original values), automated medians are strongly correlated with manual medians, indicating that relative differences among samples are well preserved, although values are consistently shifted toward larger measurements. In the bottom row (centered values), predicted medians were adjusted by subtracting the median residual for each descriptor to remove the global scaling offset. After centering, automated and manual medians show near-perfect agreement, demonstrating that the primary discrepancy reflects a consistent global bias rather than distortion of relative differences among samples.

Finally, we performed a direct comparison between predicted and manual measurements (Figure 5; Figure 3). Under these conditions, predicted and manual measurements show near-linear agreement. Across all descriptors, individual measurements cluster tightly around the identity line, and correlation coefficients exceed 0.97. This demonstrates that when the segmentation model correctly identifies and delineates an organism, the derived morphometric measurements closely match those obtained from manual annotations.

**Figure 5:**
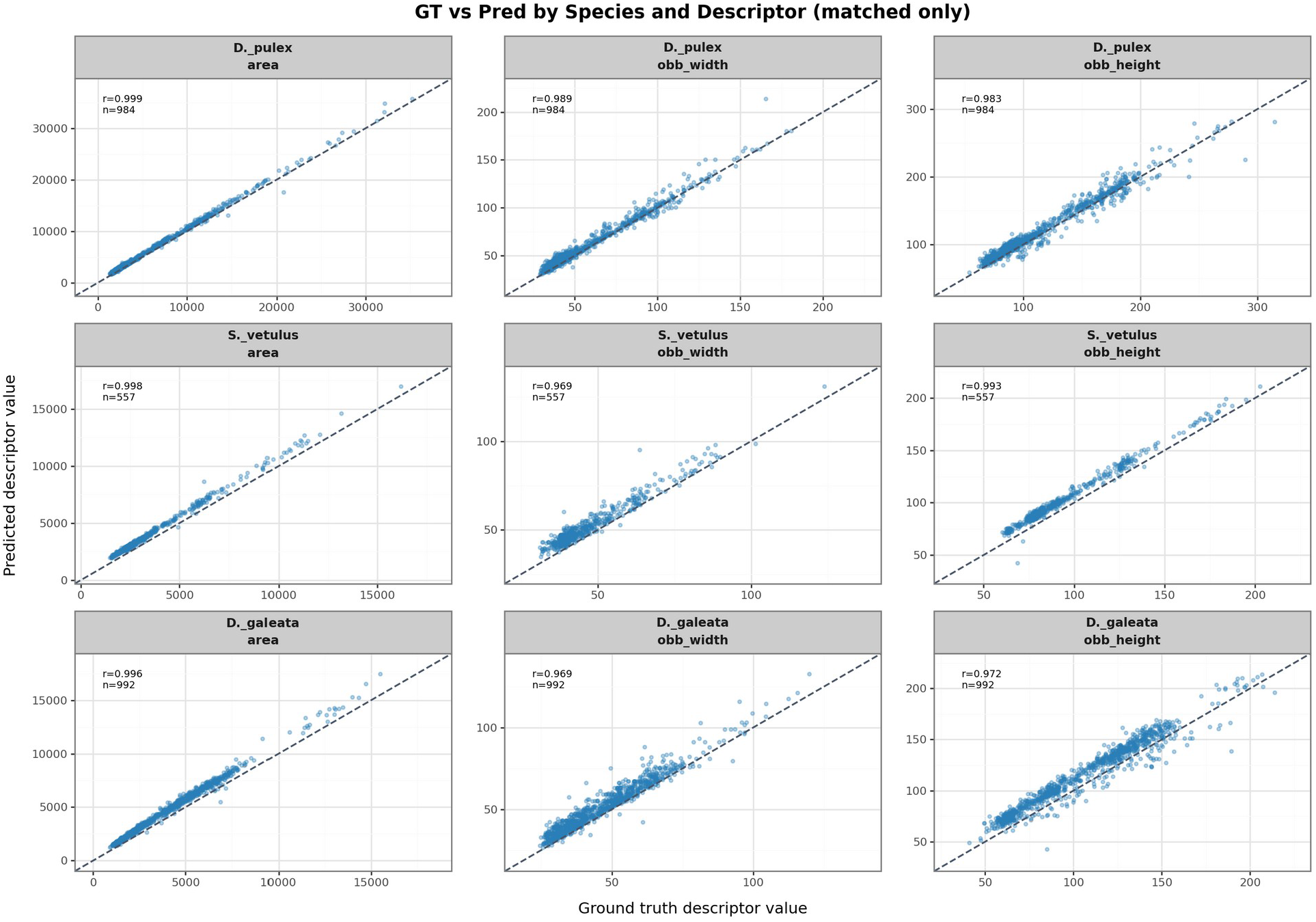
Comparison of predicted and ground-truth morphometric measurements for matched individuals across species and descriptors (IoU 0.7). Each panel shows predicted descriptor values plotted against manually derived ground-truth values for area, oriented bounding-box width (OBB width), and oriented bounding-box height (OBB height) in D. pulex, D. galeata, and S. vetulus. Points represent individual detections corresponding to the same biological organism. The dashed line indicates the identity relationship (1:1). Across all descriptors and species, predicted measurements show near-linear agreement with manual measurements (r = 0.975–0.999), indicating that when organisms are correctly detected and segmented, the derived morphometric descriptors closely match those obtained from manual annotations.

To determine whether these differences primarily reflected a stable systematic offset rather than distortion of relative trait structure, we additionally examined centered comparisons. In these analyses, predicted values were shifted by the median residual for each descriptor, so that the remaining deviations represent residual disagreement after removal of the global bias. After centering, sample-level agreement remained strong and the mean residual bias was effectively zero for all descriptors. Residual errors also remained low, with MAE) of 98.3 pixels for area, 1.42 pixels for OBB width, and 1.37 pixels for OBB height (Figure ; row: Centered). Likewise, matched-only ground-truth versus predicted descriptor plots showed strong concordance across species and descriptors, with points clustering closely around the 1:1 line and panel-wise correlations ranging from r = 0.969 to 0.999 (SFigure 3). Together, these centered results indicate that the dominant discrepancy between automated and manual measurements is a consistent scaling offset, whereas the relative ordering among samples and the internal structure of the trait distributions are preserved.

### 3.2 Robustness of Sample Median Estimates

To quantify how sampling depth affects the reliability of trait summaries, we performed a repeated subsampling analysis for each species-descriptor combination. We simulated manual trait-based sampling by randomly drawing subsamples of size n = 10, 30, and 50 individuals from the ground-truth (GT) data. For each sample size, 10,000 independent subsamples were generated, and the median trait value was calculated for each draw. Subsample error was defined as the absolute percent deviation from the full GT median. Model error was computed the same way, using the median of the full prediction set relative to the full GT median. For each subsample replicate, we then tested whether the model-derived median was closer to the GT median than the subsampled median. This allowed us to estimate, for each species, descriptor, and sample size, the proportion of cases in which the model outperformed manual subsampling. Notably, this comparison was performed on the original prediction distributions, so the model medians retained their systematic bias. The analysis therefore asks whether a model-derived summary can still be preferable to manual subsampling even when that bias is present. Overall, model-derived medians were competitive primarily at the lowest sampling depth. For *D. pulex*, the model outperformed subsampling in 53-62% of cases at n = 10 across descriptors, but this declined to 19-29% at n = 50 (Figure 6). A similar pattern was observed for *D. galeata*, where the model outperformed subsampling in 43-62% of cases at n = 10, but only 13-28% at n = 50. In contrast, for *S. vetulus*, model performance gains were limited, with the model outperforming subsampling in only 8-17% of cases at n = 10 and approaching zero at larger sample sizes (Figure 6).

**Figure 6:**
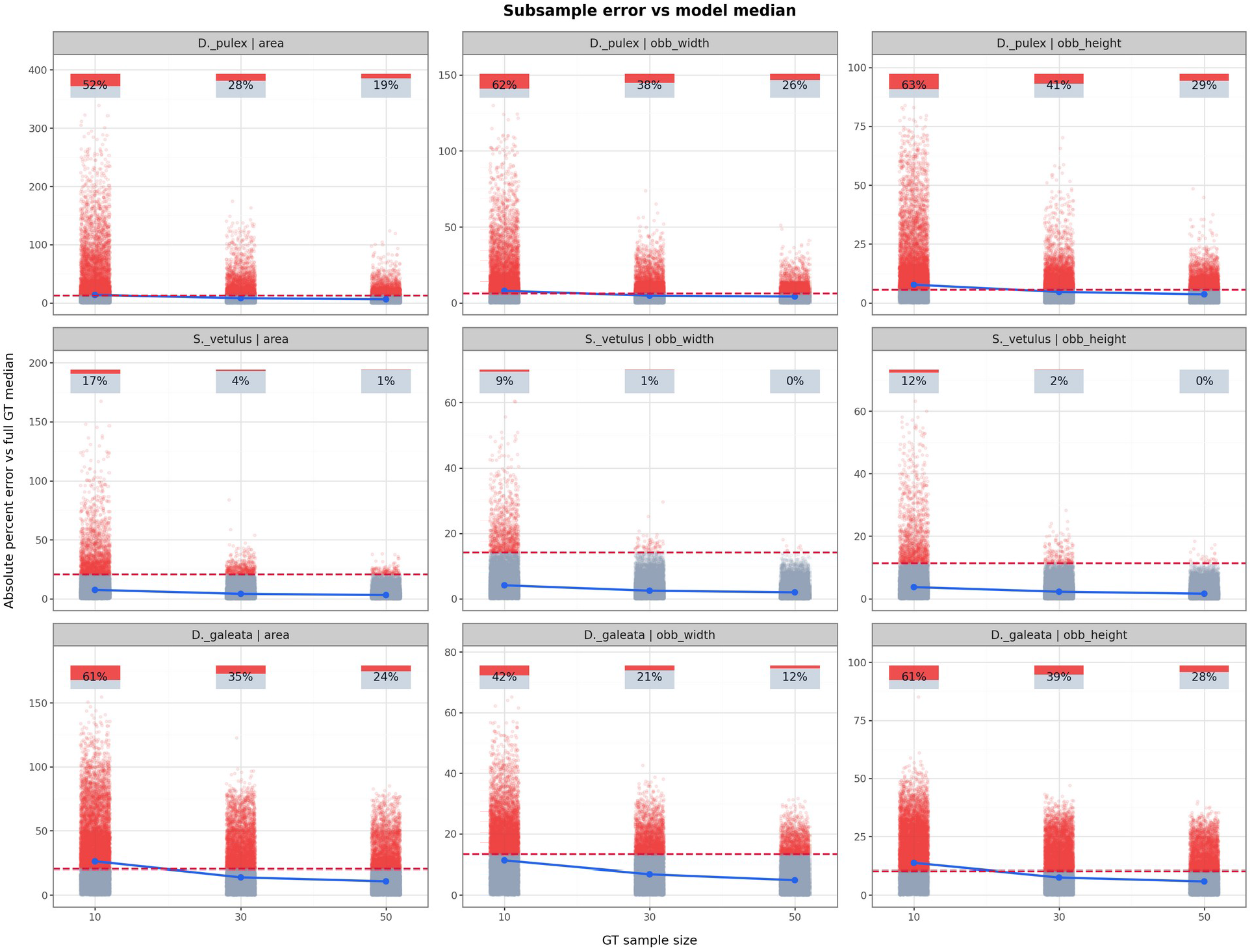
individual points represent the errors of the 10,000 GT subsamples, the blue line indicates the median subsample error across replicates, the dashed red horizontal line indicates the error of the full model median, and the percentage bar above each sample size reports the fraction of subsamples for which the model was more accurate than the subsample. To test whether model-based summaries remain useful despite systematic prediction bias, this analysis was performed on the uncalibrated prediction distributions; the same framework can also be applied to the scale-calibrated predictions to evaluate how descriptor-wise calibration changes the comparison.

### 3.3 Computational Practicality and Scalability

In addition to measurement accuracy, we evaluated the computational practicality of the proposed workflow. The complete image analysis pipeline was executed on a standard laptop computer (Intel Core i5 processor with 12 CPU cores and 15 GB RAM), without the use of specialized hardware such as dedicated GPUs. Processing a full high-resolution scan required approximately two minutes, including image chunking, instance segmentation, descriptor extraction, and result aggregation. This runtime demonstrates that the workflow can be applied routinely using widely available computing resources.

## 4. Discussion

Automated extraction of morphometric traits from high-resolution plankton images represents an important step toward upscaling trait-based ecological analyses. In aquatic ecosystems, body size is a central functional trait governing metabolic rates, grazing dynamics, and trophic transfer efficiency (Cyr & Downing, 1988; Lampert, 1987; Peters & Downing, 1984; Woodward et al., 2010). In addition, shifts in size structure often reflect ecological processes such as predation pressure, resource limitation, and environmental stress. Traditional approaches to quantifying these traits rely on manual microscopy and measurement of a relatively small number of individuals per sample (Ha & Hanazato, 2009; Milbrink & Bengtsson, 1991; Pantel et al., 2015; Van Doorslaer et al., 2010). Although these methods have produced foundational ecological insights, they are inherently time-consuming and difficult to scale, which limits the statistical power of trait-based analyses and constrains the resolution at which population-level phenotypic variation can be characterized (Culverhouse et al., 2014; Lindenmayer & Likens, 2010).

The present workflow addresses this limitation by combining instance segmentation with automated morphometric trait extraction from polygon outlines. Our assessment indicates that automated measurements preserve the structure of trait distributions and maintain strong agreement with manual measurements at the sample level. In our case, the primary discrepancy between automated and manual measurements was a consistent positive bias in absolute values. Bland–Altman analyses indicated that this bias represents a multiplicative scaling offset rather than a distortion of the underlying distribution structure. After removal of this global offset, residual differences between automated and manual measurements were small, suggesting that relative differences among samples and treatments are preserved. These findings become particularly relevant when considering realistic sampling constraints. In ecological practice, trait measurements are often based on relatively small numbers of manually measured individuals, which introduces substantial sampling variability. Our subsampling analysis (Figure 6) directly illustrates this effect. While automated estimates may exhibit a higher absolute error than many individual subsamples, the distribution of subsampling error is wide, and a substantial proportion of manual estimates deviate more strongly from the true population median. In particular, higher percentiles of the subsampling error distribution (e.g., 75th and 90th percentiles) reveal that a non-negligible fraction of manual estimates perform substantially worse than the automated approach, highlighting the intrinsic variability associated with low sampling depth. In this context, the consistency of automated measurements can outweigh their systematic bias. In other words, even though the model slightly overestimates absolute values, it does so in a systematic way, whereas manual subsampling is more sensitive to stochastic variation, especially at the low sample sizes that are typically obtained in studies (Pantel et al., 2015; Van Doorslaer et al., 2010). As sampling depth increases, this advantage naturally diminishes, as manual estimates converge toward the true population median. However, in typical use cases where large-scale manual measurements are impractical, the ability to derive stable sample-level summaries from many automatically detected individuals may be more important than achieving perfectly unbiased individual measurements.

Automated measurement also substantially increases effective sampling resolution. In addition to increasing the number of individuals analyzed per sample, this automation also enables higher sampling frequency. Because manual measurement is time-intensive, sampling is often limited to relatively coarse temporal intervals (e.g., monthly). Automated analysis substantially reduces processing time, making it feasible to increase sampling frequency (e.g., weekly or even higher resolution). This improvement allows finer-scale temporal dynamics in trait distributions to be captured, which would otherwise remain unresolved. Together, increased sampling depth and frequency enhance the statistical robustness and ecological interpretability of trait-based analyses, addressing a key limitation of traditional plankton measurement approaches (Milbrink & Bengtsson, 1991; Orenstein et al., 2022).

From a biological perspective, the usefulness of automated image analysis frameworks for ecological research largely depends on the type of information contained in their outputs. Detection-based approaches such as Slicing-Aided Hyper Inference (SAHI; (Adiwijaya et al., 2024) primarily generate bounding-box predictions around detected objects. While these outputs are effective for counting organisms, bounding boxes do not capture the true outline of organisms and therefore limit the extraction of biologically meaningful morphometric descriptors. Segmentation-based frameworks address this limitation by representing individuals through complete object contours. For example, the flatbug framework developed for terrestrial arthropod monitoring applies instance segmentation to extract organism outlines from ecological images, supporting automated biodiversity monitoring and improved size estimation (Svenning et al., 2026). However, its primary output remains segmented images of organisms rather than quantitative trait measurements, and the biological information derived from these outputs is therefore largely limited to presence, counts, or approximate size estimates. In contrast, the HiReS approach focuses on the ecological information contained in segmentation outputs by converting reconstructed organism outlines into geometric descriptors such as area, body dimensions, and shape metrics. These descriptors correspond to morphometric traits that are directly relevant to ecological processes, enabling automated characterization of trait distributions within plankton populations and communities.

Finally, automated processing also improves reproducibility in ecological monitoring workflows. Manual measurements are inherently susceptible to observer bias and differences in measurement strategy, which can introduce inconsistencies across datasets and observers (Culverhouse et al., 2014; Lindenmayer & Likens, 2010). Automated pipelines standardize measurement procedures and ensure that traits are extracted using consistent computational rules. Because the workflow can be executed on standard computing hardware and does not require specialized infrastructure, it is accessible to a wide range of laboratories and monitoring programs.

Several limitations should nevertheless be acknowledged. First, the accuracy of the workflow depends on the quality of the underlying segmentation model, and new datasets may require additional training or validation to achieve comparable performance. Second, automated measurements presented here showed a consistent positive bias relative to manual annotations, indicating that absolute descriptor values derived from automated segmentations may be systematically larger than manually derived measurements. This offset likely reflects differences in how organism boundaries are delineated rather than random measurement error. A plausible contributing factor is the inclusion of illumination-induced halos around organisms, which may lead the model to slightly overestimate true boundaries (SFigure 2).

Overall, the HiReS framework can enable a shift toward full-sample, high-resolution image analysis in zooplankton research. It offers a practical alternative to manual measurement or subsampling methods, by providing consistent and detailed trait information that enables analyses that would be otherwise prohibitively time-consuming. As machine learning is increasingly integrated into ecological research, the ability to extract reliable, individual-level measurements from large samples has the potential to support more detailed experiments, strengthen trait-based ecological inference, and improve comparability across studies and sampling campaigns.

## Declarations

## Acknowledgments

This project was supported by GK-UHH-01 Rapid Adaptive Change (Landesforschungsförderung Hamburg). We would like to thank Lyn Westphal for linguistic help.

## Authors Contributions

**SM, ST, SAJD**, and **KAO** conceived the study. **SM** developed the HiReS library, implemented the software, performed the analyses, prepared the figures, and designed the methodological framework. **ST, SAJD**, developed and optimized the scanning procedure. **ST** and **SAJD, KAO** contributed to the ecological interpretation and provided conceptual input throughout the study. **KAO** supervised the project. **SM** wrote the first draft of the manuscript. All authors contributed to revising the manuscript and approved the final version.

## Ethical Approval

This is not applicable.

## Consent to Participate

This is not applicable.

## Consent to Publish

This is not applicable.

## Data Availability

All data supporting the findings of this study are publicly available at Zenodo under the following DOI: 10.5281/zenodo.15719995.

## Supplemtary Figures

**SFigure 1:**
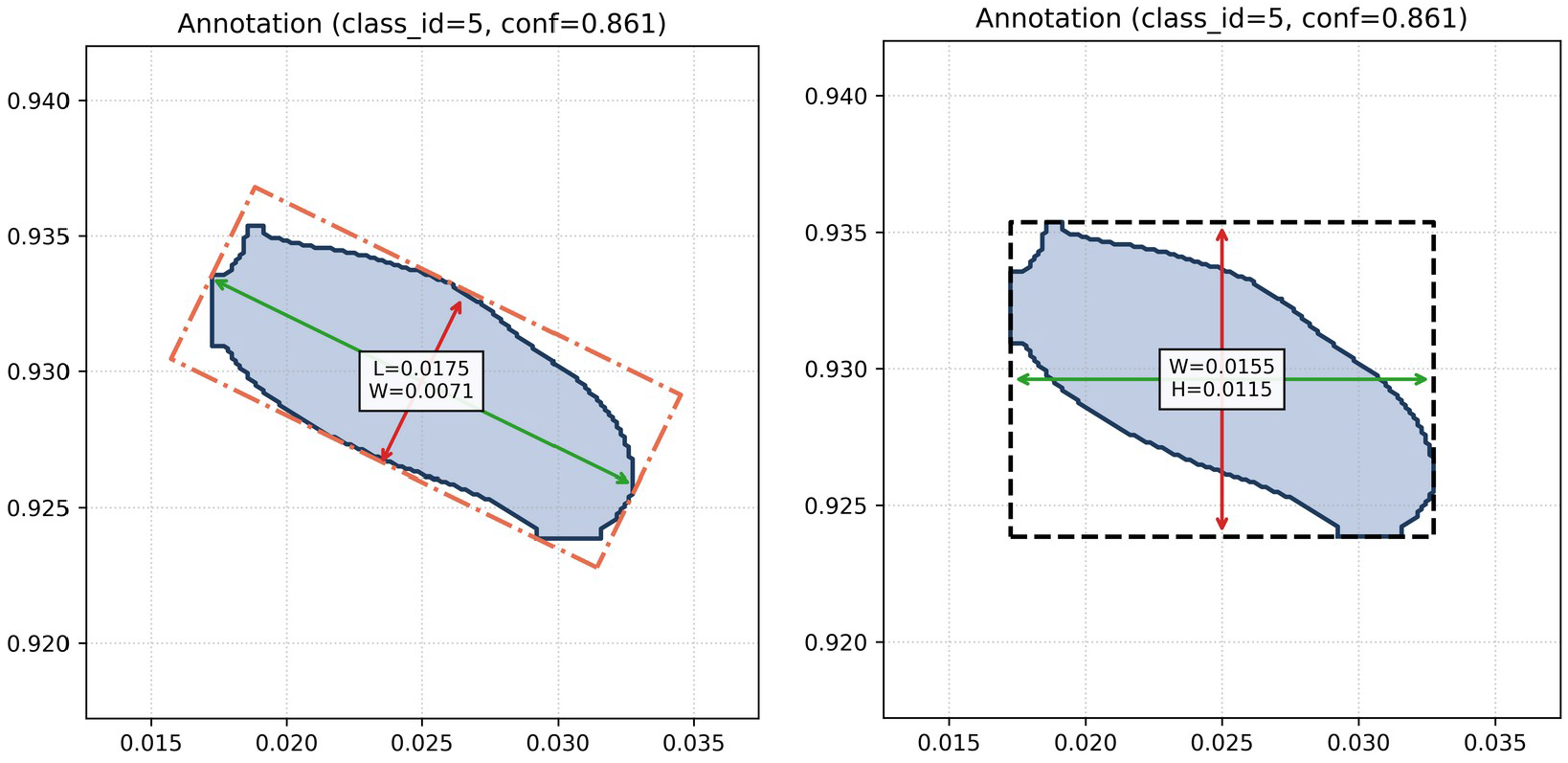
Comparison of axis-aligned and oriented bounding box–based shape measurements for a segmented object. Right: Axis-aligned bounding box (black dashed) enclosing the polygonal segmentation (blue), with width (W, green arrow) and height (H, red arrow) measured along the image axes. Left: Oriented bounding box (orange dashed) aligned with the object’s principal orientation, with length (L, green arrow) and width (W, red arrow) measured along the major and minor axes of the object. Values are reported in normalized image coordinates. The oriented bounding box provides a rotation-invariant estimate of object dimensions compared to the axis-aligned bounding box.

**SFigure 2:**
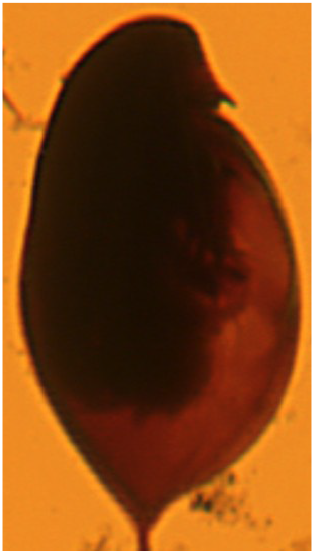
Representative example showing a halo effect caused by scanner illumination around the Daphnia, which is partially included in the predicted segmentation mask and contributes to boundary overestimation.

**SFigure 3:**
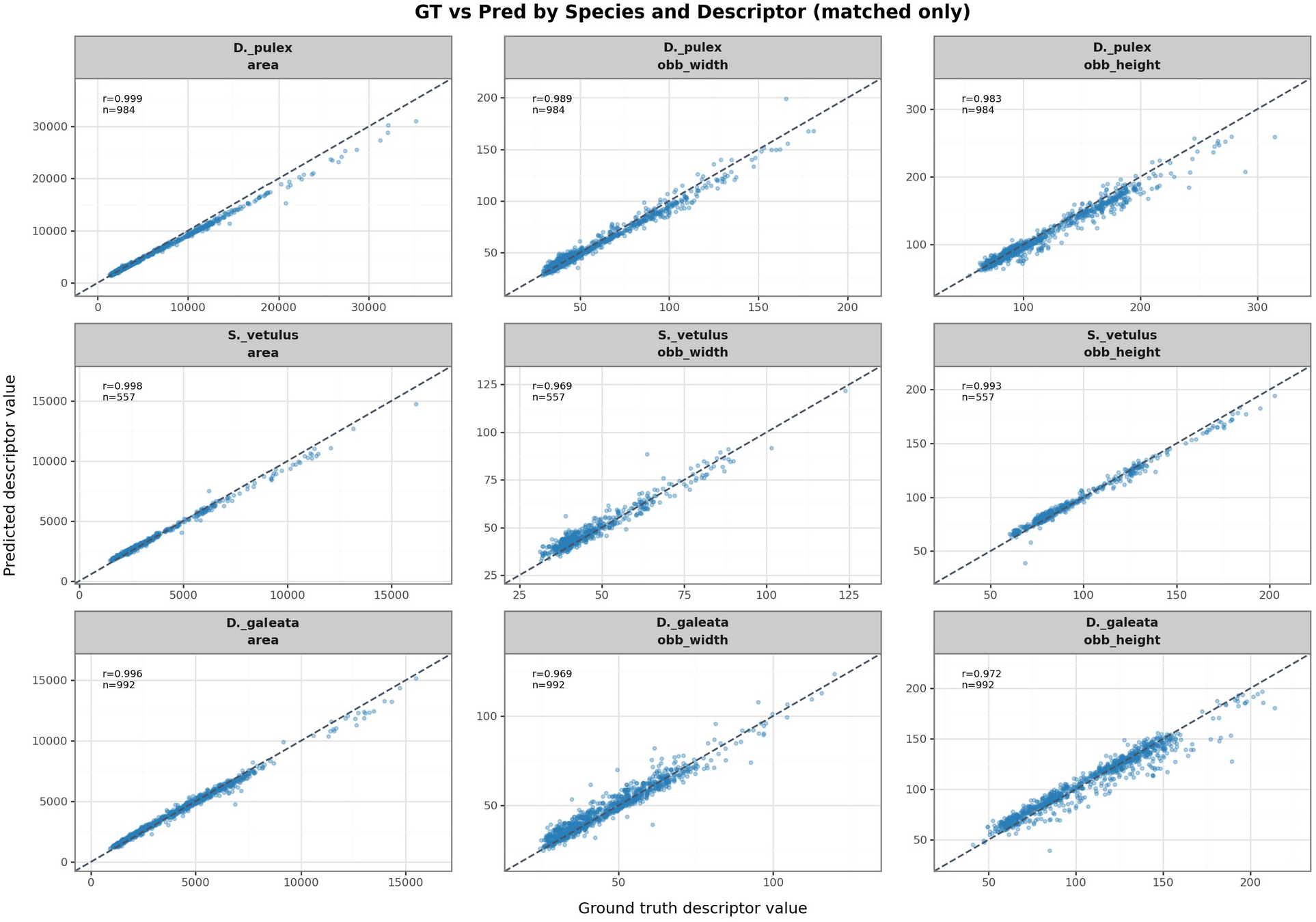
Comparison of centered predicted and ground-truth morphometric measurements for matched individuals across species and descriptors (IoU 0.7). Each panel shows prediction-derived descriptor values plotted against manually measured ground-truth values for area, oriented bounding-box width (OBB width), and oriented bounding-box height (OBB height) in D. pulex, D. galeata, and S. vetulus. Predicted values were centered by subtracting the median residual for each descriptor to remove the global scaling offset observed in the raw measurements. Points represent individual detections corresponding to the same biological organism, and the dashed line indicates the identity relationship (1:1). After centering, predicted and manual measurements show near-linear agreement across all descriptors and species, demonstrating that residual differences between automated and manual measurements are small once the systematic bias is removed.

